# Inferring CTCF binding patterns and anchored loops across human tissues and cell types

**DOI:** 10.1101/2022.06.15.496356

**Authors:** Hang Xu, Xianfu Yi, Wei Wang, Xinlei Chu, Shijie Zhang, Xiaobao Dong, Zhao Wang, Jianhua Wang, Yao Zhou, Ke Zhao, Hongcheng Yao, Zheng Nan, Junwen Wang, Dariusz Plewczynski, Pak Chung Sham, Kexin Chen, Dandan Huang, Mulin Jun Li

## Abstract

CCCTC-binding factor (CTCF) is a transcription regulator which is involved in many cellular processes. How CTCF recognizes DNA sequence to exert chromosome barrier or enhancer blocking effects remains to be fully interrogated. Despite many computational tools were developed to predict CTCF-mediated loops qualitatively or quantitatively, few could specially evaluate the regulatory potential of DNA sequence at CTCF binding sites (CBSs) and how it affects chromatin loop formation. Here, we developed a deep learning model, DeepAnchor, to precisely characterize the binding patterns for different types of CBSs. By incorporating base-wise genomic/epigenomic features, we revealed distinct chromatin and sequence features for CTCF-mediated insulation and looping at a high resolution, such as two sequence motifs flanking the core CTCF motif at loop-associated CBSs. Besides, we leveraged the predicted anchor score to optimize the loop extrusion model and achieved the best performance in predicting CTCF-anchored loops. We established a compendium of context-specific CTCF-anchored loops across 52 human tissue/cell types and found that genomic disruption of CTCF-anchored loops may represent a general causal mechanism of disease pathogenesis. These computational models, together with the established resource, could facilitate the mechanistic research on how the CTCF-mediated *cis*-regulatory elements (CREs) shapes context-specific gene regulation in cell development and disease progression.

## Introduction

Variable interactions between enhancer and target gene have been systematically profiled in explaining tissue/cell type-specific transcriptional regulation (Consortium, 2012; Consortium et al., 2020), but the mechanisms underlying the precise wiring between genes and the enhancers from varied distances remain unclear. As the main insulator-related transcription factor (TF) discovered in vertebrates, CTCF assists cohesin to form chromatin loops, which is believed to be the fundamental biophysical basis for distal gene regulation (Ali et al., 2016; Braccioli and de Wit, 2019; Gabriele et al., 2022; Ghirlando and Felsenfeld, 2016; Nichols and Corces, 2015; Ong and Corces, 2014; Phillips and Corces, 2009). According to the loop extrusion model (Davidson et al., 2019; Kim et al., 2019), cohesin dimers load on DNA at NIPBL binding sites and slide along with DNA until reaching the anchors, which are usually bound by specific cofactors, such as CTCF, YY1, and ERα (Beagan et al., 2017; He et al., 2014; Sanborn et al., 2015). Chromatin loops mediated by YY1 and ERα are found to form enhancer-promoter interactions, while CTCF-anchored loops generally enclose the formers (Beagan et al., 2017), assembling genome units that are named insulated neighborhoods (INs). Many lines of evidence also suggest that these structures constitute the mechanistic underpinnings of higher-order chromosome organizations (Beagan and Phillips-Cremins, 2020; Hnisz et al., 2016), such as topologically associating domains (TADs) (Dixon et al., 2012) (Dixon et al., 2012; Nora et al., 2012). Enhancers tend to regulate genes within the same IN/TAD while the regulation across different INs/TADs is more likely to be prevented (Dowen et al., 2014; Islam et al., 2023). Therefore, characterizing CTCF binding patterns and their regulatory consequences will greatly facilitate the analyses of fine-grained 3D genome regulation.

CTCF and its associated *cis*-regulatory elements (CTCF-mediated CREs) play a critical role in controlling tissue/cell type-specific gene expression by maintaining chromatin domain boundaries or blocking enhancer activities, however, CTCF has been revealed as a versatile TF that plays several other roles in different scenarios, including gene activation, transcriptional repression, pausing, and alternative splicing (Filippova et al., 1996; Guo et al., 2012; Lobanenkov et al., 1990; Oh et al., 2021; Shukla et al., 2011; Vostrov and Quitschke, 1997). Therefore, it is necessary to distinguish insulator or looping-related CTCF binding sites (CBSs) from those with other functions. There are a lot of works that analyze the functional heterogeneity of CTCF in the nucleus, which revealed that the binding motif of CTCF could be the key to understand the behavior of CTCF at specific site (Huang et al., 2021b; Nakahashi et al., 2013; Ribeiro-Dos-Santos et al., 2022). Studies have uncovered that insulation potency heavily depends on the number of CBSs in tandem and an upstream flanking sequence at the core CTCF motif (Huang et al., 2021b). Besides, in the formation of CTCF/cohesin chromatin loops, the CBSs at loop anchors tend to be arranged in convergent orientation (Guo et al., 2015). However, these studies are either unable to count the feature patterns of individual CBSs or limited to given genomic loci. Besides, a large number of computational tools were developed to qualitatively or quantitatively predict CTCF-mediated loops (Cao et al., 2021; Ibn-Salem and Andrade-Navarro, 2019; Kai et al., 2018; Kuang and Wang, 2021; Lv et al., 2021; Matthews and Waxman, 2018; Oti et al., 2016; Wang et al., 2021; Xi and Beer, 2021; Zhang et al., 2018), but few could specially evaluate the regulatory potential of DNA sequence at CBS and how it affects the loop formation. Moreover, it remains elusive to see which factors or combinations of features determine the specificity among different types of functional CTCF binding events, and whether such information could contribute the prediction of CTCF-anchored loops across the whole genome.

Here, we develop an interpretable deep learning model, DeepAnchor, to ask what types of genomic/epigenomic features determine a CBS to be insulator/cohesin/loop-associated. By leveraging large-scale base-wise genomic and epigenomic features, we uncovered several critical patterns for CTCF-mediated insulation and looping at a high resolution. We embedded the predicted anchor score to the previous loop extrusion model (Xi and Beer, 2021), and found the optimized model LoopAnchor outperforms the existing methods for predicting CTCF-anchored loops. Furthermore, for the first time, we established a landscape of tissue/cell type-specific CTCF-anchored loops across 52 human tissue/cell types and applied them to interpret disease-causal variants. This method together with the compiled resources will facilitate the mechanistic research on the dynamic regulation of 3D chromatin during cell development and disease progression.

## Results

### DeepAnchor enables high-confidence CTCF binding patterns characterization

The key feature of DeepAnchor is the implementation of a deep learning model that leverages base-wise features for CTCF-medicated CREs detection. Previous studies usually analysed the sequence and chromatin status of CBS in particular tissue/cell types with a one-dimensional feature vector, in which regional feature values are averaged for each CBS regardless of heterogeneous signals across the locus (Kai et al., 2018; Zhang et al., 2018). However, a true CTCF-medicated CRE differs from other CBSs in fine-grained architecture (Huang et al., 2021b; Lee et al., 2022; Ong and Corces, 2014), which raises new requirements for feature characterization before modelling. One straightforward solution is to profile a CTCF binding event using high-resolution features surrounding the CBS. In our implementation, for each CBS, 44 quantitative base-wise genomic/epigenomic features within the ± 500bp region were obtained from CADD annotation (Rentzsch et al., 2019), and DNA sequence within ±500bp of the CBS was also extracted and represented by one-hot encoding (Table S1). Thus, by concatenating large-scale annotations with converted DNA features at the base-wise level, a feature matrix with a size of 1000×48 was generated for each CBS (Figure 1A top left).

**Figure 1.**
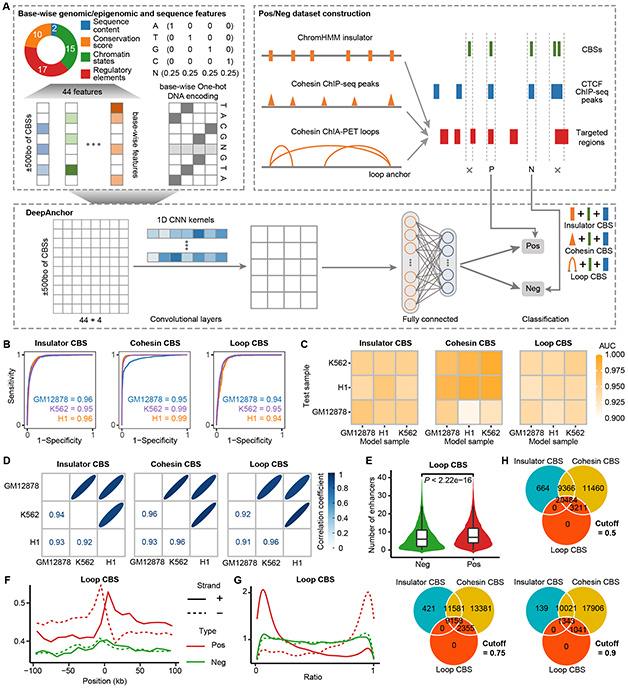
Structure and performance of DeepAnchor model. A. Schematic view of DeepAnchor model. Base-wise chromatin features and DNA sequences are extracted for all candidate CBSs by motif scanning. Positive (Pos) and negative (Neg) dataset is constructed by considering both CTCF ChIP-seq peaks and targeted chromosome intervals, including ChromHMM insulator-associated CBS (insulator CBS), cohesin ChIP-seq signal-associated CBS (cohesin CBS), and cohesin ChIA-PET loop-associated CBS (loop CBS). 1D-CNN model is then used to train a classifier for distinguishing positive CBSs (CTCF-mediated CREs) versus other ones. Probability of CBSs being insulator/cohesin/loop-associated can be calculated and used as DeepAnchor score for downstream analyses. **Related terminologies**: **Insulator**, a enhancer blocker or a barrier between heterochromatin and euchromatin; **Chromatin loop**, during the interphase of cell, the condensed chromatin form 3D structure within cell nucleus. The basic loop-like structure is called chromatin loop; **Loop anchor**, given a chromatin loop detected by ChIA-PET, we call the endpoints of loop on chromosome as loop anchor; **Insulator/cohesin/loop CBS**; by using different targeted regions, we get different P/N dataset and train different DeepAnchor models. The CBS predicted by different models will be named by particular targeted region. B. Cross-validation ROC curves based on GM12878, K562 and H1-hESC datasets, respectively, among three types of CBSs. C. Cross-sample ROC curves for DeepAnchor models on different cell types, among three types of CBSs. D. Correlation between DeepAnchor scores for three cell type-specific models, among three types of CBSs. E. Comparison of number of enhancers around CBSs between predicted Pos and Neg CTCF-mediated CREs in loop CBS model. F. Strand-oriented asymmetric pattern of enhancer enrichment at predicted Pos and Neg CTCF-mediated CREs in loop CBS model. G. Position and strand preference of TAD boundary enrichment by measuring the distance between each CBS and the 5’ end of TAD it belongs at predicted Pos and Neg CTCF-mediated CREs in loop CBS model. H. The intersection of the predicted CTCF-mediated CREs among three CBS types at different thresholds.

We modelled three major forms of CBSs across the whole human genome, including insulator-associated CBS (insulator CBS), cohesin-associated CBS (cohesin CBS) and loop-associated CBS (loop CBS). These CTCF-medicated CREs share common CTCF binding motif but could display varied regulatory functions. To prepare high-quality ground truth for different types of CBS learning, we required that each selected positive site should contain 1) putative CBS by motif scanning; 2) observed CBS by CTCF ChIP-seq at given tissue/cell type. For insulator CBS, we intersected the selected sites with genomic segments showing ChromHMM insulator state (15 states model). For cohesin CBS, we overlapped the selected sites with cohesin ChIP-seq in matched cell type. For loop CBS, we required the selected sites located in cohesin loop anchor by ChIA-PET at same cell type (Figure 1A top right). We also randomly selected equal number of negative samples from the whole CBS pool. To avoid ambiguous results, multiple CBSs shared the same CTCF/cohesin ChIP-seq peak(s) or the same cohesin loop anchor(s) are excluded (see Methods). Including base-wise features of the region that spans the selected CBSs will largely expand the feature volume and scale, which will introduce a flood of information that is usually beyond the ability of classical machine learning algorithms. For example, regions with 1000bp can incorporate 1000×48=48,000 features if a base-wise value is applied. To address this challenge, we applied a deep convolutional neural network (CNN) to extract important patterns from the feature matrixes, which greatly reduces the calculation burden and could increase prediction performance. After training a classifier on a positive and negative dataset, the DeepAnchor framework returns the probability of a CBS region belonging to a true CTCF-mediated CRE, which is called the DeepAnchor score in the downstream analyses (Figure 1A bottom, see Methods).

We trained DeepAnchor model using high-quality CTCF/RAD21 ChIP-seq and RAD21 ChIA-PET data on the GM12878, K562, and H1-hESC cell lines respectively, and tested the repeatability of DeepAnchor by cross-validation (see Methods). To avoid the potential influence of chromosome-specific confounding factors, we divided the train-test-valid set by splitting CBSs into subsets each on different chromosomes. That is, we selected chr1-16 as the training set, chr17-18 as the validation set, and others as the test set. The DeepAnchor models trained on different types of CBS samples all received high area under the ROC curve (AUC, 0.94-0.99) via five-fold cross-validation across the three cell lines (Figure 1B). To demonstrate that the DeepAnchor scores derived from different samples are comparable, we performed cross-cell type-validation across the three cell lines. Notably, among different types of CBSs, the DeepAnchor models exhibited consistently high cross-sample prediction performance on all testing pairs (all AUCs > 0.9, Figure 1C) and the DeepAnchor scores are highly correlated among the three cell type-specific models (Pearson’s r = 0.91-0.96) (Figure 1D), suggesting CTCF-mediated CREs share conservative features in different contexts and the feasibility to predict genome-wide functional CBSs at the organismal level.

We set a cut-off to define positive and negative CTCF-mediated CREs by finding the points on the ROC curve with the largest Youden’s J statistic. The optimal cut-point was obtained when the DeepAnchor score is close to 0.5 for different models, where 86%-95% positive and 85%-88% negative samples in training datasets are correctly assigned. With the GM12878 DeepAnchor models and the optimal cut-off, we generated positive and negative CTCF-mediated CREs genome-widely. Using the FANTOM5 enhancer dataset, we found that positive CBSs colocalize with more enhancers within ±100kb regions than negative CBSs on average (P-value < 2.22e-16, Mann Whitney U Test) (Figure 1E and Figure S1A). Besides, by counting the number of enhancers in every 10kb bin within ±100kb of CTCF-mediated CREs, we noted that there is a strand-oriented asymmetric pattern for positive CBSs, but not negative ones, particularly for loop CBSs and cohesin CBSs (Figure 1F and Figure S1B). Consistent with previous findings (Clarkson et al., 2019), the density of CTCF-mediated CREs around the TAD boundaries also shows strand bias, in which positive CBSs at plus-strand are usually enriched at the left TAD boundary, while those at minus-strand are aggregated at right TAD boundary. However, no such patterns were observed for negative CBSs (Figure 1G and Figure S1C). We also calculated the intersection of the predicted CTCF-mediated CREs (at different thresholds) among three CBS types. We found that all loop CBSs and most of insulator CBSs are cohesin-related, which indicates that CBS bound by cohesin could has other functions than forming chromatin loop or insulation. When increasing the cutoff of DeepAnchor score, the overlap between the three types of CBSs decreased a lot, and only the overlap between insulator CBSs and cohesin CBSs have slight changes (Figure 1H). Together, these results demonstrated that our DeepAnchor model can accurately capture high-confidence CTCF-mediated CREs across the whole genome.

### DeepAnchor reveals distinct base-wise chromatin and sequence features for CTCF-mediated CREs

Although DeepAnchor leverages a deep learning structure to extract high-level configurations from large-scale features, it also raises challenges in analysing feature importance and its biological insight. Base-wise representation of various genomic/epigenomic features in the DeepAnchor model makes it possible to depict a comprehensive picture of the patterns of CTCF-mediated CREs. To visualize underlying relationships among these features, we first clustered the feature scores upon the training dataset and found these features could be generally partitioned into three major subsets, including transcription-associated, conservation-associated, and chromatin state-associated feature sets (Figure S2). We used SHapley Additive exPlanations (SHAP) (Lundberg et al., 2020), a game-theoretic approach to explaining the output of DeepAnchor. SHAP returns the Shapley value matrix with the same alignment as the input feature matrix, which shows the contribution of each feature at each position. To inspect the overall contribution of each feature, we calculated the mean Shapley values at all positions and then summarized the mean values (Figure 2A). Overall, feature of CTCF binding evidence at open chromatin (EncOCctcfPval) greatly contribute to the model. Shapley values of G/T/A/C are also top-ranked, which implies the importance of DNA sequence. As expected, open chromatin features (e.g., EncOCFairePVal, EncOCC) display higher importance than other features (Figure 2A and Figure S3). Interestingly, nucleosome positioning evidence (EncNucleo) displayed more contribution in loop CBS model, suggesting the CTCF footprinting pattern among nucleosomes could determine the formulation of CTCF-mediated chromatin loop. Besides, a conservation feature, the neutral evolution score defined by GERP++ (GerpN) also showed weak preference in loop CBS model (Figure 2A). Taken together, these results indicate that some informative features may capture the unique and important patterns that can distinguish CTCF-mediated CREs from other types of CTCF binding events.

**Figure 2.**
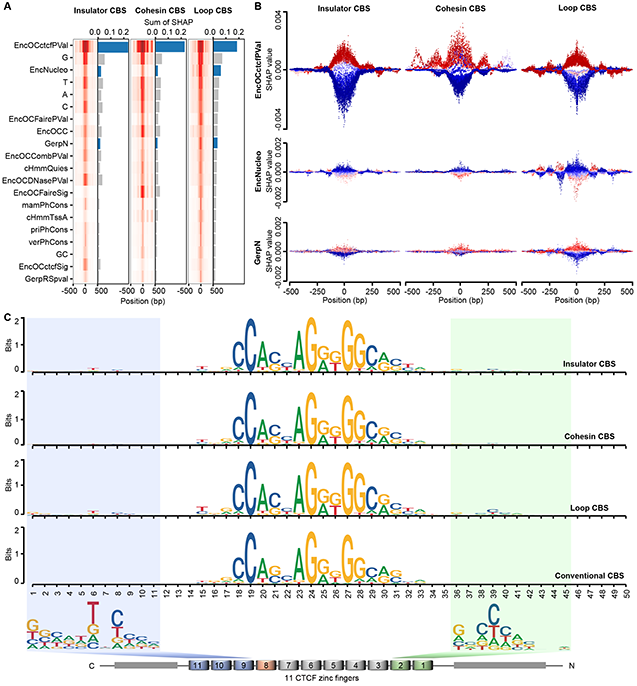
Base-wise analysis of sequence and chromatin patterns across different types of CTCF binding. A. Feature importance analysis of top 20 features at base-wise level among three types of CBSs. Heatmap: Average absolute feature Shapley values of each position at ±500bp of CBS. Barplot: Summation of absolute feature Shapley values across ±500bp of CBS. B. Base-wise Shapley value distribution for three representative features across different types of CBSs. EncOCctcfPval: P-value (PHRED-scale) of CTCF evidence for open chromatin; EncNucleo: Maximum of ENCODE Nucleosome position track score; GerpN: Neutral evolution score defined by GERP++. Please refer to more feature descriptions from supplementary table. C. Comparison of DNA motifs associated with positive CBSs among three types of CBSs in this study and a commonly used conventional CBS.

To explore the base-wise schema of CTCF-mediated CREs, for several important features, we investigated the distribution of Shapley values derived from the DeepAnchor model at every position of CBS and its surrounding region (±500bp) upon the testing dataset with randomly sampled CBSs. For example, CTCF binding intensity and its occupancy (EncOCctcfPval) display a periodical pattern at cohesin/loop CBSs instead of insulator CBSs, in which higher binding intensity of central CBS heavily contributes to positive CBS discrimination but the Shapley values flip at around 150bp far away from central CBS (Figure 2B top). Interestingly, we observed that, only at loop CBS, the Shapley values of nucleosome positioning signature (EncNucleo) are reciprocally distributed with those of CTCF binding intensity (Figure 2B middle), and the nucleosome positioning level is anti-correlated with its Shapley value (Figure S4A), indicating that CTCF at loop anchor could enlarge and protect the linked DNA between its two neighboring nucleosomes to facilitate the loop formation. Such base-wise visualization of feature contribution also greatly supports the previous finding that the binding of CTCF provides an anchor point for positioning nucleosomes thus resulting in an array of well-positioned nucleosomes flanking sites at particular CBSs (Cuddapah et al., 2009; Fu et al., 2008). We also found that the Shapley values of the conservation feature (GerpN) showed similar distribution as those of CTCF binding features, especially in loop CBSs (Figure 2B bottom). In addition, consistent with previous results (Luan et al., 2021), an investigation of base-wise Shapley value distribution upon positive or negative training datasets demonstrated that CTCF-mediated CREs usually demonstrated stronger binding intensity than other types of CBSs (Figure S4B).

Previous CTCF multivalency studies revealed that CTCF reads sequence diversity through combinatorial clustering of its 11 zinc fingers (ZFs) (Nakahashi et al., 2013; Rhee and Pugh, 2011), yet the sequence determinants of CTCF-mediated insulation remain elusive. By applying MEME motif discovery to all predicted positive and negative CBSs, we revealed two additional weak motifs flanking the core CTCF motif only at positive loop and insulator CBSs (Figure 2C). In agreement with a sensitive insulator reporter assay (Huang et al., 2021b), the upstream motif separated from the core sequence by 5 or 6 bp is recognized by ZFs 9-11 and may contribute to the directionality of CTCF binding to CBSs (Yin et al., 2017). Interestingly, the downstream motif next to the core sequence is different from previous findings in terms of distance and sequence context (Nakahashi et al., 2013). Nonetheless, this downstream sequence associated with CTCF N terminus and ZFs 1-2 has been documented to stabilize cohesin engagement (Li et al., 2020). Thus, the distinct sequence features together with multiple lines of existing evidence for CTCF binding patterns further demonstrate the fidelity of the DeepAnchor model.

### Incorporating DeepAnchor score into loop extrusion model improves the performance of CTCF-anchored loop prediction

Although DeepAnchor models the insulative and looping potential of genome-wide CBSs at the organismal level, accurate prediction of context-specific CTCF-anchored loop will be essential in understanding 3D chromatin interactions and fine-grained gene regulation. By assuming that looping interactions found in the CTCF ChIA-PET experiments are due to the blocking and localization of cohesin at CBSs, a simple but effective loop extrusion model (LEM) was previously developed (Xi and Beer, 2021). This quantitative model evaluates CTCF-anchored loop formation with only four features, including CTCF-binding intensity, CTCF motif orientation, the distance between CTCF-binding events, and loop competition. Albeit the acquired specificity, we reasoned that incorporating the loop-associated DeepAnchor score into LEM could further improve the performance of CTCF-anchored loop prediction. To this end, by considering the looping potential of CTCF binding (anchor score predicted by loop-associated DeepAnchor model), we weighed the probability of CTCF binding in the original LEM model and reached a new implementation, called LoopAnchor (see Methods).

We compared the performance of LoopAnchor with five prevalent algorithms for CTCF-mediated interaction prediction, including Naïve and Oti methods (Oti et al., 2016), CTCF-MP (Zhang et al., 2018), Lollipop (Kai et al., 2018), and LEM (Xi and Beer, 2021). To ensure a fair benchmark, all supervised models were trained on GM12878 RAD21 ChIA-PET data and tested on K562 RAD21 ChIA-PET data. We found that LoopAnchor outperforms other state-of-art methods in terms of both ROC-AUC (0.936) and PR-AUC (0.921) (Figure 3A), indicating the effectiveness of our model optimization. Besides, the correlation analysis between the predicted loop intensities and the observed ChIA-PET signals on K562 showed that LoopAnchor receives the highest correlation (Pearson’s r = 0.653) among the six tools (Figure 3B), which also demonstrates the superiority of LoopAnchor for quantitative measurement of CTCF-anchored loop.

**Figure 3.**
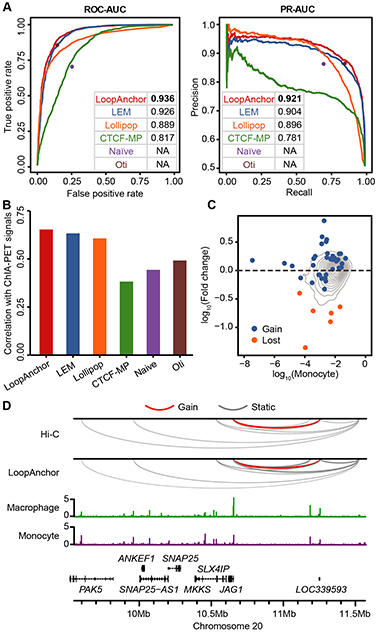
Performance evaluation of LoopAnchor for CTCF-anchored loop prediction. A. ROC curves, Precision-Recall curves and associated AUCs among LoopAnchor and five state-of-the-art methods. All supervised models, such as LoopAnchor, LEM, Lollipop and CTCF-MP were trained on GM12878 RAD21 ChIA-PET data and independently tested on K562 RAD21 ChIA-PET data. B. Correlation between predicted scores and real RAD21 ChIA-PET loop intensity on K562. C. Gain and lost loops from monocyte to macrophage differentiation by comparing LoopAnchor prediction with Hi-C observation. log10(Fold change) is the transformed fold change of predicted loop intensity for specific loop between macrophage and monocyte; log10(Monocyte) is the transformed Hi-C loop score observed in monocyte; blue dot is lost Hi-C loop; orange dot is gained Hi-C loop. D. Example of loops predicted by LoopAnchor at *JAG1* locus. For Hi-C loops, line color is used to distinguish gained and static loops. For LoopAnchor, linewidth represents the predicted loop intensity and the loop with fold change of intensity >3 was marked with red color.

To investigate whether our LoopAnchor mode could capture the dynamic loop changes during cell development, we applied our LoopAnchor model to human monocytes activation data with paired *in situ* Hi-C and CTCF ChIP-seq before and after exposure to PMA (Phanstiel et al., 2017). We found our LoopAnchor model can accurately predict 25/34 gained loops and all six lost CTCF-anchored loops (Figure 3C). For example, on the *JAG1* locus at chromosome 20:9547732-11575097, there are eight loops detected by Hi-C, among which one gained loop is identified only at differentiated macrophages (Figure 3D). Consistently, LoopAnchor not only detected all unchanged loops but also accurately predicted the gained loop (Figure 3D, see Methods). Notably, in this genomic region, the difference of loop intensity between monocyte and macrophage cells predicted by LoopAnchor is highly correlated with the loop score changes measured from Hi-C data (Pearson’s r = 0.796), indicating that LoopAnchor can quantitatively capture the local loop structure changes at the dynamic cellular process.

### A landscape of CTCF-anchored loops across 52 human tissue/cell types

Given the simplicity and gained performance of the LoopAnchor model, we sought to leverage existing large-scale CTCF ChIP-seq data to learn a global picture of CTCF-mediated chromatin interactions across different human tissue/cell types. We first uniformly processed 740 CTCF ChIP-seq datasets collected from ENCODE (Consortium et al., 2020), CistromDB (Zheng et al., 2019), and ChIP-Atlas (Oki et al., 2018) based on the ENCODE TF ChIP-seq processing pipeline (Table S2, see Methods). According to data quality control, peak calling filtering, and redundancy removing, we totally retrieved 168 high-quality CTCT ChIP-seq datasets, relating to 113 normal and 55 cancer samples across 32 tissues and 20 anatomical cell types (Figure 4A and Table S3, see Methods). Each biosample contains around 100,000 peaks (mean = 104,691, median = 96,296, Figure S5A). We applied LoopAnchor and LEM to detect genome-wide CTCF-mediated loops for each biosample respectively, and we found that nearly 20,000 loops were detected per biosample (mean = 16,255 and median = 16,637 for LoopAnchor, mean = 17,826 and median = 18,252 for LEM, Figure S5B and S5C).

**Figure 4.**
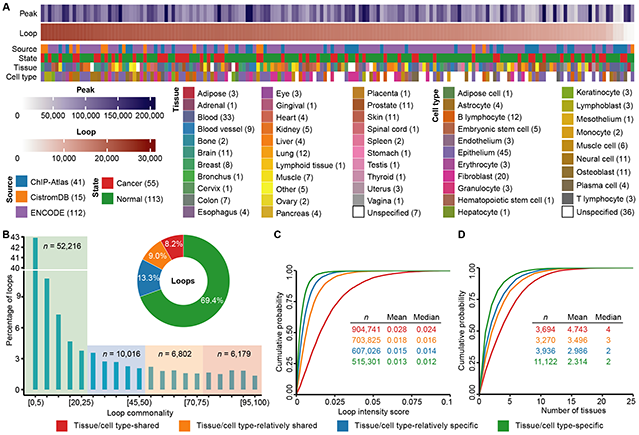
The landscape of predicted CTCF-anchored loops across 32 human tissues and 20 cell types. A. Overview of predicted CTCF-anchored loops across 168 biosamples (columns) and biological conditions (rows). B. Classification of CTCF-anchored loops according to their shared patterns. All predicted loops were classified into four categories, including tissue/cell type-specific, tissue/cell type-relatively specific, tissue/cell type-relatively shared, and tissue/cell type-shared. C. Comparison of loop intensity score for loops in four categories. The cumulative probability was calculated and the two-tailed Mann-Whitney U test was used to test the significance. D. Validation of tissue distribution for loops across four categories using predicted chromatin loops by Peakachu for 42 human tissue/cell types. The cumulative probability was calculated and the two-tailed Mann-Whitney U test was used to test the significance.

Based on the shared pattern of detected loops across 168 biosamples, we first classified all LoopAnchor loops into four types: (1) 8.2% (*n* = 6,179) tissue/cell type-shared loops in more than 75% biosamples; (2) 9.0% (*n* = 6,802) tissue/cell type-relatively shared loops in more than 50% but less than 75% of biosamples; (3) 13.3% (*n* = 10,016) tissue/cell type-relatively specific loops in more than 25% but less than 50% of biosamples; (4) 69.4% (*n* = 52,216) tissue/cell type-specific loops in less than 25% of biosamples (Figure 4B, see Methods). Based on the same strategy, the LEM loops were also classified into these four types (Figure S6A). Compared to LEM, LoopAnchor can detect more shared loops but less specific loops globally (8.2% vs. 3.9% for tissue/cell type-shared loops, and 69.4% vs. 80.3% for tissue/cell type-specific loops, Figure 4B and Figure S6A), suggesting LoopAnchor could capture more conservative CTCF-anchored loops across different tissue/cell types. By investigating the distribution of LoopAnchor-predicted loop intensity scores, we found a significant difference among the four loop categories (P-value < 2.22e-16, two-tailed Mann-Whitney U test, Figure 4C), in which the tissue/cell type-shared loops received the highest scores and the tissue/cell type-specific loops got the lowest scores. A similar pattern was also observed for the loop intensity score derived from the LEM method (Figure S6B). To evaluate the tissue/cell type specificity of classified CTCF-mediated loops, we leveraged the predicted chromatin loops by Peakachu for 42 human tissue/cell types (Salameh et al., 2020) and compared the number of associated tissue/cell types among different categories. We found that the tissue/cell type distribution of the LoopAnchor-predicted loops also showed significant differences among the four loop categories (P-value < 2.22e-16, two-tailed Mann-Whitney U test, Figure 4D). As expected, the number of associated tissue/cell types gradually decreased from the tissue/cell type-shared group (mean = 4, median = 4.743) to the tissue/cell type-specific group (mean = 2, median = 2.314), indicating the effectiveness of our loop classification. A similar trend was also identified for the detected loops from the LEM method (Figure S6C). Together, we established a compendium of tissue/cell type commonality and specificity for CTCF-anchored loops across large-scale human tissue/cell types. All predicted loops for 168 selected biosamples and all 764 biosamples can be visualized and compared as separated tracks at UCSC Track Data Hubs (https://genome.ucsc.edu/cgi-bin/hgHubConnect) by entering customized hub URLs https://raw.githubusercontent.com/mulinlab/LoopAnchor/master/hubs_landscape.txt or https://raw.githubusercontent.com/mulinlab/LoopAnchor/master/hubs_all.txt, respectively.

### Tissue/cell type-specific loop anchors are highly enriched at disease-causal loci

Multiple lines of evidence have supported that CTCF/cohesin binding sites are highly mutated in cancers (Fang et al., 2020; Guo et al., 2018; Katainen et al., 2015) or constantly shaped via evolutionary selection (Kentepozidou et al., 2020; Sadowski et al., 2019). However, whether context-specific CTCF-mediated looping and associated loop anchors are more likely linked to disease-causal genomic loci has not been systematically tested. By collecting 12,738 causal variants for 54 blood-related autoimmune diseases from genome-wide association studies (GWASs) (Wang et al., 2020) (Table S4) and sampling matched control variants, we evaluated the genome-wide enrichment of autoimmune disease-causal variants on the CTCF-anchored loops predicted by LoopAnchor in normal tissues with more than three biosamples (see Methods). Notably, the autoimmune disease-causal variants were significantly enriched in the loop anchors in blood tissue (mean P-value = 0.00357) compared to other tissues (mean P-value ranges from 0.14465 (colon) to 0.0055 (prostate), Figure 5A). As the majority of disease-causal regulatory variants show tissue/cell type specificity in phenotypically relevant contexts, these results further support that context-dependent loop information could better interpret GWAS disease-causal variants for complex diseases.

**Figure 5.**
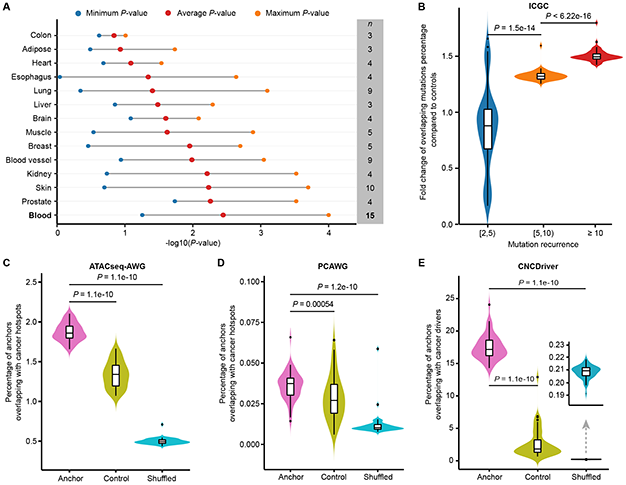
Disease-causal variants and somatic hotspots enrichment. A. Enrichment significance distribution for autoimmune disease-causal variants among different normal tissues. The tissues were ordered by their average P-values. B. Comparison of fold change distribution among different somatic mutation recurrence categories. The two-tailed Mann-Whitney U test was used to test the significance. C-E. Comparison of overlapped percentage with somatic hotspots using different datasets from ATACseq-AWG (C), PCAWG (D), and CNCDriver (E). The paired two-tailed Mann-Whitney U test was used to test the significance.

To evaluate the relevance of CTCF-mediated loop anchors in the cancer mutation process, we collected somatic mutations from ICGC whole genome mutation aggregation (Consortium, 2020) and extracted recurrent mutations in the non-coding regions (see Methods). By comparing with permuted non-recurrent mutations, we found that mutations with higher recurrence obtain greater enrichment in CTCF-mediated loop anchors for cancer biosamples (two-tailed Mann-Whitney U test, Figure 5B). Besides, based on the whole-genome cancer mutation hotspots collected from three pan-cancer whole-genome analyses, including ATACseq-AWG (Corces et al., 2018), PCAWG (Rheinbay et al., 2020), and CNCDriver (Liu et al., 2019), we tested whether the mutation hotspots are more frequently occurred in CTCF-mediated loop anchors than other CTCF binding loci or shuffled non-coding genomic regions (see Methods). We uncovered consistent results among different datasets, in which the loop anchors were highly enriched with cancer mutation hotspots (two-tailed Mann-Whitney U test, Figure 5C-F). These results suggest genomic disruption of CTCF-anchored loops may thus represent a general causal mechanism of disease pathogenesis.

## Discussion

The multiple critical functions of CTCF in the nucleus stimulate constant phenotypic and mechanistic research in the field, yet how CTCF recognizes insulators to exert chromosome barrier or enhancer blocking effects remains to be interrogated. In this study, we developed a novel computational model to accurately predict CTCF-mediated CREs and CTCF-anchored loops genome-widely. By incorporating large-scale base-wise genomic and epigenomic features within a deep learning model, we revealed several distinct chromatin and sequence features for CTCF-mediated insulation at a high resolution. Importantly, we found two additional sequence motifs flanking the core CTCF motif at positive loop and insulator-associated CBSs. Besides, we leveraged the predicted insulator score to optimize the previous loop extrusion model and achieved better performance in predicting CTCF-anchored loops. Based on the model, for the first time, we established a compendium of tissue/cell type-specific and -shared CTCF-anchored loops across 52 human tissue/cell types. Finally, we showed that tissue/cell type-specific loop anchors are highly enriched at disease-causal loci. Therefore, our results deepen the understanding of CTCF-mediated insulation and loop formation. The new method together with the compiled resource also provides useful approaches for studying the dynamic regulation of 3D chromatin during cell differentiation and disease progression.

Although many computational models that can predict CTCF-mediated loops with varied sensitivity and specificity (Cao et al., 2021; Deng et al., 2022; Ibn-Salem and Andrade-Navarro, 2019; Kai et al., 2018; Kuang and Wang, 2021; Lv et al., 2021; Matthews and Waxman, 2018; Oti et al., 2016; Wang et al., 2021; Xi and Beer, 2021; Zhang et al., 2018), they usually learn from CTCF ChIA-PET data and barely evaluate the regulatory potential and loop attribute among different types of CBS. Here, we applied cohesin ChIA-PET/ChIP-seq data, ChromHMM-predicted insulator, and strict CTCF binding evidence to specifically analyze CTCF binding patterns at three types of CTCF-mediated CREs. This provides a unique tool for characterizing CTCF binding consequence and loop formation through the loop extrusion mechanism. Notably, the limited availability and varied library construction quality of CTCF/cohesin ChIA-PET restrict the training of context-specific models broadly, however, our cross-cell type comparisons showed a high agreement among DeepAnchor models trained from different ChIA-PET data, indicating the discrimination of true CTCF-mediated CREs may rely on several conservative features revealed by our interpretable deep learning model, such as higher binding intensity, well-positioned nucleosomes as well as two unique motifs flanking the central CBS.

By introducing the DeepAnchor score into a loop competition and extrusion model (Xi and Beer, 2021), we observed improved performance in predicting CTCF-anchored loops and demonstrated that the new LoopAnchor method can achieve better quantitative estimation. However, in both LEM and LoopAnchor, the quantitative predictions only measure the contribution of each component independently. For example, the insulation potential of CBS and loop competition are dependent processes based on dynamic chromatin context (Davidson and Peters, 2021), which motivates a future optimization direction. Besides, given existing methods without providing CTCF-associated anchor prediction results, we cannot evaluate whether incorporating other anchor score would achieve better performance. Instead, we performed systematic comparisons with five prevalent algorithms for CTCF-mediated loop prediction. Clearly, our method is limited to inferring specific CREs and loops bound by CTCF, but recent studies indicated that some TFs could function as novel architectural proteins in regulating genome organization (Yi et al., 2021), such as YY1 (Weintraub et al., 2017), ZNF143 (Bailey et al., 2015), MAZ (Ortabozkoyun et al., 2022), BHLHE40 (Hu et al., 2020), CTCFL (Debruyne et al., 2019), etc., and some of which are independent of CTCF binding. Our model can be easily extended to train classifiers to extract base-wise features for binding patterns of these architectural proteins, given the targeted regions and negative controls. Thus, predicting full spectrum anchor sites and their associated looping events across the genome deserve further in-depth investigation.

The landscape of CTCF-anchored loops established by applying LoopAnchor to 168 uniformly processed human CTCF ChIP-seq biosamples could be a useful resource in 3D genome and regulatory genomics studies. Since anchor scores were estimated at the organism level, LoopAnchor still only requires CTCF binding profiles to accurately predict CTCF-anchored loops. This will greatly simplifies the application, particularly for studying dynamic 3D CTCF code during cell development and disease progression. In addition, the compiled tissue/cell type-shared and -specific loops could facilitate the interpretation of disease-causal variants identified by GWAS and somatic non-coding driver mutations in cancers.

## Methods

### Training dataset preparation

We prepared three types of target regions from high quality functional genomics data of three Tier 1 ENCODE cell lines (GM12878, K562, and H1-hESC), including insulators estimated by ChromHMM (Ernst et al., 2011) (https://genome.ucsc.edu/cgi-bin/hgTrackUi?db=hg19&g=wgEncodeBroadHmm), cohesin (RAD21) ChIP-seq narrow peaks (ENCODE: ENCSR000AKB, ENCSR000BPJ, ENCSR000BNH), and cohesin (RAD21) ChIA-PET loops (ENCODE: ENCSR752QCX, ENCSR000CAC, ENCSR543YTV) (Grubert et al., 2020). For ChromHMM insulator segments and RAD21 ChIP-seq narrow peaks, we merged the intervals to avoid overlapping. For RAD21 ChIA-PET data, we preprocessed them with ChIA-PET2 using default parameters (Li et al., 2017). Inter-chromosomal interactions and interactions with less than 2 PETs were filtered out, and the anchor of loops were extracted and merged based on position to get anchor list without overlapped regions. Then, three types of CBSs were prepared by intersecting target intervals with candidate CBSs by motif scanning as well as CTCF ChIP-seq narrow peaks (ENCODE: ENCSR000AKB, ENCSR000BPJ, ENCSR000BNH), including insulator CBS, cohesin CBS, and loop CBS. Thus, for each cell type, the candidate CBSs colocalized with both CTCF ChIP-seq peaks and targeted regions were selected as positive set, while others constituted the pool of negative set. To avoid biases, we removed the CBSs shared the same target region from positive regions, because we cannot distinguish which one is the true positive one.

### Feature preparation

DNA binding motif of CTCF (MA0139.1) was downloaded from JASPAR2020 (Fornes et al., 2019). We used FIMO of MEME Suite (v5.1.1) (Grant et al., 2011) to scan the CTCF binding motif across the whole human genome (GRCh37/hg19), resulting in 110,059 CTCF binding sites with P-value < 1e-5. According to the topic relevance, we selected 44 base-wise genomic/epigenomic features from the CADD (v1.4) annotation database (https://cadd.gs.washington.edu/static/ReleaseNotes_CADD_v1.4.pdf) (Kircher et al., 2014; Rentzsch et al., 2019). For each CBS, feature values of ±500kb surrounding region were extracted and stored together to form a feature matrix. All features were scaled by a min-max scaling algorithm. To reduce the influence of extreme high/low values, we extracted 0.1 and 99.9 percentile as the min and max values. Besides, we got the DNA sequence of the ±500kb region of the CBS center using BEDTools (Quinlan and Hall, 2010). A, T, C, G were then transferred into numerical value with one-hot encoding: A (1,0,0,0), T (0,1,0,0), G (0,0,1,0), C (0,0,0,1). Finally, by concatenating large-scale annotations with converted DNA features at the base-wise level, a feature matrix with a size of 1000×48 was generated for each CBS.

### DeepAnchor model and fitting

Here we described the statistical model of DeepAnchor to identify insulator/cohesin/loop-associated CBSs. For each CBS *n* = 1, …, *N*, we extracted genomic/epigenomic features and DNA sequence features at a base-wise level for the ±500bp region of the CBS center. By concatenating features and one-hot encoding sequence, a two-dimensional 1000×48 signal matrix *x*_*n*_ is generated for each CBS *n*. To extract feature patterns from the signal matrix, DeepAnchor uses 1D convolutional kernels to process the signal matrix. For each layer *l*, the forward propagation from layer *l* − 1 to *l* is expressed as,

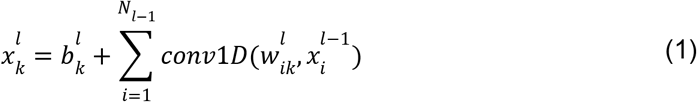

Where *N*_*l*_ is the number of neurons at layer *l*, 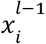 and 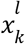 are the *i^th^* neuron at layer *l* − 1 and *k^th^* neuron at layer *l*, 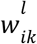 and 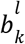 are the kernel and the scalar bias from layer *l* − 1 to *layer l*. DeepAnchor implements two 1D-CNN layers, each followed by a max-pooling layer and a drop-out layer. After 1D-CNN layers, all the outputs are fully connected. Three fully connected layers convert the outputs of 1D-CNN layers to lower dimensions and finally train a classifier with sigmoid activation,

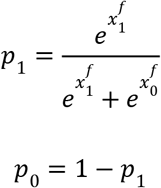

where *x^f^* is the result of fully connected layers which contain only two elements for positive training and negative target respectively. *p*_1_ is the probability that CBS *n* belongs to a positive CBS. The DeepAnchor model was implemented with TensorFlow framework. To avoid overfitting, we extracted a balanced negative set by randomly selecting samples from the pool of negative set. Finally, positive and negative datasets were divided into train, test, and valid set according to chromosomes. Samples on chr1-16 were selected as the training set, chr17-18 as validation set and the others as test set. We considered *p*_1_ and *p*_2_ as the possibility the CBS to be a true target-associated CTCF or not, and we calculated binary cross entropy as the loss of model. Input data were divided into small batches with batch size of 50. The model was trained for 20 epochs, but early stopping was enabled when the validation loss does not decrease any more.

### Evaluation of DeepAnchor model

We performed five-fold cross-validation to evaluate the DeepAnchor model and plotted ROC curves and corresponding AUC. We set a cut-off to define positive and negative CBSs by finding the points on the ROC curve with the largest Youden’s J statistic. Same with the training procedure on GM12878, DeepAnchor models were trained for each cell type and the models were also validated by all test sets belonging to three cell types. After training the DeepAnchor model, we applied it to all potential CBSs across the whole human genome. DeepAnchor will calculate a score ranging from [0, 1], with a larger value meaning more likely to be a positive CTCF-mediated CREs. Human enhancer annotation was downloaded from the FANTOM5 database (Lizio et al., 2015) (https://fantom.gsc.riken.jp/5/datafiles/latest/extra/Enhancers/). According to Youden’s J statistic, CBSs with anchor scores > 0.5 are selected as positive CTCF-mediated CREs, while negative CTCF-mediated CREs are those with anchor scores < 0.5. We also investigated the performance of DeepAnchor models with different cutoffs. We counted the number of enhancers for positive/negative insulators if the distance between them is smaller than 100kb. We tested the differences of involved enhancers between positive and negative CBSs using the Mann Whitney U Test. The estimated TAD data was derived from the GM12878 Hi-C assay (Dixon et al., 2012), in which there are 4,386 TADs and the median length of TADs is 520kb. To test whether CBSs have preference for a position or not, we counted the distance between each CBS and the 5’ end of TAD it belongs. Since TADs have variable lengths, we normalized the distance by counting the ratio concerning the total length of the TAD. Therefore, the normalized distance of CBSs will be the percentage of relative position within TAD.

### Base-wise feature contribution analyses

We used SHapley Additive exPlanations (SHAP) (Lundberg et al., 2020) to interpret the output of DeepAnchor. Since the input of the DeepAnchor model is a 48×1000 feature matrix, SHAP returns the Shapley value matrix with the same alignment as the input feature matrix. We leveraged the shap package (Lundberg et al., 2018) to load the well-trained DeepAnchor model and used the training dataset as background examples to receive an expectation over. Then an explainer was generated based on the model and background dataset. The explainer was then applied to predict the Shapley values for every position via testing datasets. Note that the training dataset and testing dataset were the same as the ones in the training DeepAnchor model. We also interpreted the feature contribution based on the mean Shapley values for all positions.

### Optimization of loop extrusion model in LoopAnchor

Enlightened by a recent loop competition and extrusion model (Xi and Beer, 2021), we integrated the loop-associated DeepAnchor score (estimated from loop CBS DeepAnchor model) into the model to predict CTCF-anchored loops. We estimated the probability of insulator-associated CTCF binding at CBS *i*:

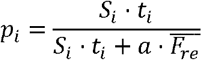

where *S*_*i*_ is the CTCF ChIP-seq signal, and *a* is a constant which have been estimated in the original paper. In this new implementation, *t*_*i*_ is the loop-associated DeepAnchor score at CBS *I*, thus, revised mean CTCF signal 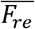 is given by:

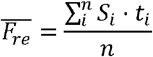

### Comparison with state-of-art methods for CTCF-mediated loop prediction

We acquired Oti code from Oti et.al (Oti et al., 2016), and the naïve method was derived from Oti code by setting the recursive round of the Oti method to 1. We used the default parameters to run Oti and Naïve methods, and loops with a score > 0 were considered positive loops while others were treated as negative ones. CTCF-MP (Zhang et al., 2018), Lollipop (Kai et al., 2018), and LEM (Xi and Beer, 2021) were installed and used on relevant cell types according to their GitHub repositories. To facilitate comparison among different methods, all methods were trained on the GM12878 cell line and tested on the K562 cell line. Balanced (Pos/Neg = 1) and unbalanced (Pos/Neg = 1/5) gold datasets were prepared respectively. ROC curve, PR curve, and corresponding AUC were then plotted and calculated. We also calculated the Pearson correlation coefficient between predicted loop intensity with real data to show the performance of all methods.

### Application of LoopAnchor for dynamic loop detection during macrophage development

We downloaded CTCF ChIP-seq data for both monocyte and macrophage cells (Phanstiel et al., 2017) (GEO: GSE96800). CTCF ChIP-seq data with replication were processed with ENCODE standard pipeline to retrieve bigWig and peak files. LoopAnchor was used to predict CTCF-anchored loops with peak files for both cell states. We also obtained cell state-specific based on Hi-C loops detected by HiCCUPS (Rao et al., 2014) (GEO: GSE63525). Because Hi-C loops are not necessary to be mediated by CTCF and cohesin, we filtered out loops without CTCF peaks in anchor regions on both sides. We mainly focused on the ‘gained’ and ‘lost’ loops from monocyte to macrophage cells by comparing the predicted loop intensity *L* of the same loop between the two cell states. The fold change of loop intensity for a specific loop between macrophage and monocyte is defined as *L*_*macro*_/*L*_*mono*_ and thresholding at |*L*_*macro*_/*L*_*mono*_ | > 3 for ‘gained’ and ‘lost’ loops.

### Landscape construction of CTCF-anchored loops across human tissue/cell types

We systematically collected CTCF ChIP-seq datasets from ENCODE (Consortium et al., 2020), CistromDB (Zheng et al., 2019), and ChIP-Atlas (Oki et al., 2018), and uniformly processed them using the ENCODE TF ChIP-seq processing pipeline (https://github.com/ENCODE-DCC/chip-seq-pipeline2). Briefly, the clean reads were mapped to the human genome (GRCh37/hg19) with BWA (v0.7.17) (Li and Durbin, 2009) followed by the post-alignment filtering, and finally, the peaks were called using SPP (v1.15) (Kharchenko et al., 2008). To select the high-quality ChIP-seq biosamples, we firstly filtered out the outliers with excessively low or high peaks. Then, all B lymphocyte biosamples prefixed with “GM” were removed except GM12878. For each mapped tissue or anatomical cell type, we dropped redundant biosamples according to Jaccard similarity clustering of called peaks and selected one biosample in each cluster with a maximal peak number. Finally, we applied LoopAnchor and LEM to detect genome-wide CTCF-mediated loops for each biosample respectively.

### Definition of tissue/cell type-sharded and -specific loops

All predicted CTCF-anchored loops were generally classified into four categories according to their shared patterns across different tissue/cell types. Specifically, loops that were shared in no less than 75% tissue/cell types were classified as tissue/cell type-shared loops. Loops that were shared by no less than 50% but less than 75% tissue/cell types were classified as tissue/cell type-relatively shared loops. Loops that were shared by no less than 25% but less than 50% tissue/cell types were classified as tissue/cell type-relatively specific loops. The remaining loops which were shared by less than 25% tissue/cell types were classified as tissue/cell type-specific loops. The public Hi-C loops detected by Peakachu (Salameh et al., 2020) for 42 tissue/cell types were downloaded from the 3D genome browser (Wang et al., 2018) (http://3dgenome.fsm.northwestern.edu/publications.html). Then, the “pairtopair” command from BEDTools (Quinlan and Hall, 2010) with parameter “-type both” was used to compare them with loops predicted by LoopAnchor.

### GWAS disease-causal variant enrichment

We collected 12,738 causal variants for 54 blood-related autoimmune diseases from CAUSALdb (Wang et al., 2020). To test the genome-wide enrichment of autoimmune disease-causal variants on the CTCF-anchored loops predicted by LoopAnchor, we first sampled the same amount of control variants 10,000 times with matched allele frequency using vSampler (Huang et al., 2021a). For each normal biosample, the predicted CTCF-anchored loops were flattened into unique anchors, and the “intersect” command from BEDTools (Quinlan and Hall, 2010) was used to perform the colocalization between variants and anchors. The enrichment P-value was calculated by counting the number of permutations that receive a higher percentage of overlapping variants than the real dataset in each biosample. This analysis was only performed on normal tissues with more than three biosamples.

### Non-coding somatic mutation and hotspot enrichment

The genome-wide somatic mutations were downloaded from ICGC Data Portal (Release 28) (Consortium, 2020). Candidate non-coding somatic mutations were retained by removing these overlapped with exons and splicing sites and were classified into three categories according to their mutation recurrence (∈[2,5), ∈[5,10), ≥10). The somatic mutations with only one recurrence were used as the background for drawing control datasets. We generated 10,000 control datasets by randomly sampling the equivalent mutations for each category. For each cancer biosample, the predicted CTCF-anchored loops were flattened into unique anchors, and the count of intersected hits between somatic mutations in each category and anchors in a particular biosample was calculated. By comparing to the average count value from the control datasets, the fold change of overlapping mutation percentage was derived. The Mann-Whitney U test was used to test the difference in fold changes among the three categories. In addition, we also curated candidate somatic mutation hotspots or driver regions from three previous catalogs, including ATACseq-AWG (Corces et al., 2018), PCAWG (Rheinbay et al., 2020), and CNCDriver (Liu et al., 2019). For each cancer biosample, the loops predicted by LoopAnchor with a score ≥ 0.01 were kept and flattened to unique anchors as the “Anchor” datasets, while the same number of non-anchor regions were sampled from remaining loops with the smaller predicted scores as “Control” datasets. By excluding real hotspots and restricting them to non-coding genomic regions, the “shuffle” command from BEDTools (Quinlan and Hall, 2010) was also used to retrieve shuffled hotspots with 100 permutations as “Shuffled” datasets. The Mann-Whitney U test was used to test the difference in the overlapped percentage of somatic hotspots among these generated datasets.

## Data availability

All predicted loops for 168 selected biosamples and all 764 biosamples can be visualized and compared as separated tracks at UCSC Track Data Hubs (https://genome.ucsc.edu/cgi-bin/hgHubConnect) by entering customized hub URLs https://raw.githubusercontent.com/mulinlab/LoopAnchor/master/hubs_landscape.txt or https://raw.githubusercontent.com/mulinlab/LoopAnchor/master/hubs_all.txt, respectively.

## Code availability

The source codes of LoopAnchor are freely available under MIT License at https://github.com/mulinlab/LoopAnchor or https://bitbucket.org/xuhang01/loopanchor.

## Contributions

M.J.L. and H.X. conceived of the project. H.X., X.Y. and D.H. performed the analyses and implemented the software. W.W., X.C., D.H., S.Z., X.D., Z.W., J.W., Y.Z., K.Z., H.Y. and N.Z. contributed the data collection and processing. J.W., K.C., D.P. and P.C.S. contributed to manuscript polishing and provided analysis suggestions. H.X., X.Y. and M.J.L. wrote the manuscript. The authors read and approved the final submission.

